# Snail plots are badges of genome assembly quality

**DOI:** 10.1101/2025.11.20.689594

**Authors:** Richard Challis, Mark Blaxter

## Abstract

Assembly quality is frequently assessed using independent measures of assembly span, contiguity, sequence composition and completeness. Among contiguity metrics, contig and scaffold N50 have perhaps gained the most traction, despite well known limitations. Several authors have suggested considering the complete N*x* curve rather than just the N50 value, but while using N90 values or considering the area under the N*x* curve with auN statistics provide more complete measures of contiguity, they share the limitation of being unsuited to direct comparison across a range of genome sizes. We introduced snail plots to provide a genome-size independent way to summarise a range of commonly used assembly metrics. Here we demonstrate that easily-learnt visual differences between snail plots allow simultaneous consideration of metrics across several key areas of assembly quality to rapidly identify high- and low-quality assemblies. We show that prominent features in snail plots of high-quality assemblies effectively highlight N50, N90 and auN contiguity statistics, but since the presentation is scaled to the longest scaffold, plots can be compared effectively across a wide range of taxa and assembly sizes. We use the core features of a snail plot to derive a proportional measure of assembly quality based on auN, adjusted for ambiguous, N, bases and scaled to the length of the longest scaffold. We show that this “snail score” value corresponds closely to a qualitative assessment of overall assembly quality from visual interpretation of a snail plot.

## Introduction

Genome assembly is the process of reconstructing the DNA sequence of an organism from available sequencing data. Higher quality assemblies are those that provide a more accurate representation of the true DNA sequence in terms of overall contiguity, sequence composition, base-level accuracy and completeness. Assessment and comparison of assemblies typically relies on a set of metrics that each provide a partial measure of one of these aspects of assembly quality.

Assembly contiguity is commonly measured using the N50 length of the constituent scaffold (or contig) sequences. N50 is the length of a scaffold such that at least 50% of the total span of the assembly is covered by scaffolds of that length or greater and its counterpart NG50 provides the same measure but relative to the expected true genome size. These values can be calculated for any percentage value, *x*, so N50/NG50 are part of a family of N*x*/NG*x* metrics. These metrics have limitations as they cannot be directly compared across assemblies of taxa with different-sized genomes (Miller et al. 2010) or with widely divergent chromosome counts, they may be biased upwards by misassembly of large scaffolds, and they do not reflect increases in contiguity for scaffolds below the N*x*/NG*x* length. Several authors have addressed these last points by considering the full N*x*/NG*x* curves for a set of assemblies (e.g. (Bradnam et al. 2013)) and defining a metric based on the sum of squared scaffold lengths divided by the sum of scaffold lengths as E-size (the expected size of a scaffold that would contain a gene chosen at random (Salzberg et al. 2012)) or auN (the area under the N*x*/NG*x* curve (Li 2020).

Species-specific differences in sequence composition, such as in mean GC proportion, can be used to identify non-target data in genome assemblies ((Kumar et al. 2013)). However, intra-genomic variation in GC proportion can be greater than inter-genomic variation due to differences between coding and non-coding regions and the presence of isochores. In some cases leading to bi- and multi-modal distributions of GC-proportion across a genome ((Salinas et al. 1988; Lander et al. 2001)). GC proportion is therefore most useful as an indication of assembly quality in combination with other sources of evidence.

Sequence accuracy can be measured in terms of the overall error rate, with large-scale biodiversity sequencing projects such as the Vertebrate Genome Project setting a threshold of Q40, or 1 error per 10,000 bases as a minimum acceptable metric (Rhie et al. 2021). Alternatively accuracy can be assessed against markers of assembly completeness using single copy orthologues.

Assembly completeness itself may be directly measured by the length of the assembly compared to the expected length of the full genome sequence. Other measures include the proportion of contigs assigned to chromosomes, and the presence of telomeric repeats at the ends of chromosomes for a full telomere-to-telomere assembly.

Completeness can also be estimated by exploration of expected gene content. The Benchmarking Using Single Copy Orthologues (BUSCO) system uses a set of genes believed to be near-universally present and in single copy in all species within a given taxonomic group (Simão et al. 2015). While there may be lineage specific variation in expected BUSCO completeness, a high score is an indication that the full genic content of a genome is included in the assembly. High rates of BUSCO duplication may indicate uncollapsed heterozygosity, while fragmentation of BUSCO genes may be a marker of low contiguity.

Full assessment of assembly quality requires consideration of a combination of these metrics and several tools are available to perform quality assessments, often incorporating additional data from a reference assembly or mapped reads to allow assessments of correctness, e.g. QUAST (Gurevich et al. 2013), LASER (Khiste and Ilie 2015), Merqury (Rhie et al. 2020), WebQUAST (Mikheenko et al. 2023) and Genome Evaluation Pipeline (Sullivan et al. 2025). Such tools include tabular and graphical outputs, but interpreting the core statistics requires adjustment of expectations according to the overall expected genome size and chromosome count. Comparing tabulated results across assemblies also implies a focus on a subset of metrics and brings issues of inconsistent formatting when comparing across different sources.

Snail plots were conceived as an assembly “badge” to provide a consistent way to convey a suite of these metrics for any assembly. Relative to tables of core metrics, snail plots convey a greater density of information, with a high “data to ink” ratio (Tufte 2001) making it easier to assess core aspects of assembly quality at a glance, use consistent formatting to reduce the cognitive load of comparing data from different sources, and are scale-independent, so expectations do not need to be adjusted based on the size of the genome under consideration. Snail plots also convey the distribution of scaffold lengths so provide similar insight to metrics such as auN. These distributions can highlight features of assemblies that may be missed by other metrics, such as a high number of very short scaffolds.

Snail plots were first introduced in the LepBase genome browser (Challis et al. 2016), for which they were implemented as a set of standalone scripts (Challis 2017). They have since been re-implemented as a core component of BlobToolKit (Challis et al. 2020), with snail plots for over 10,000 assemblies available on the public viewer at blobtoolkit.genomehubs.org/view. Snail plot assembly badges also appear as main figures in numerous genome assembly papers, including over 1,300 genome notes from the Darwin Tree of Life project [e.g. (Mead et al. 2020), (Aunin et al. 2021)] and are presented in a modified form on WormBase ParaSite (Howe et al. 2017).

Here we describe the most recent implementation of snail plots and provide best practice guidance for configuring, presenting and interpreting the plots both as an assembly quality inference tool and to offer insight into high level assembly organisation. We also suggest a new metric based on area under N that provides a single value for assembly quality that can be compared across assemblies of different sizes and correlates closely with visual indicators of assembly quality observed in snail plots.

## Methods

### Implementation

Interactive snail plots are available as part of the BlobToolKit Viewer (Challis et al. 2020). Snail plots may also be generated from any BlobToolKit dataset, either available as a local directory, or remotely hosted via the BlobToolkit API using the BlobTK command `blobtk plot`. BlobTK is a suite of tools that provides high-performance functions for processes commonly used in BlobToolKit and other GenomeHubs tools.

BlobTK is implemented in Rust for optimal performance when processing assemblies with their associated reads and read alignments. While these considerations are less directly relevant to the `blobtk plot` command, using the same codebase allows for efficient processing of JSON BlobDir files via serde (https://github.com/serde-rs/serde) and serde_json (https://github.com/serde-rs/json) and SVG generation with resvg (https://github.com/linebender/resvg). The Command line interface is implemented with clap (https://github.com/clap-rs/clap) and python bindings to `blobtk plot` are implemented with maturin (https://github.com/PyO3/maturin) to allow the plotting functionality to be imported into python code.

### Making a snail plot

#### Data preparation

Snail plots can be generated from an assembly FASTA file and BUSCO full table TSV file, for which any relevant BUSCO lineage may be used, but convention is to use the most specific lineage available. The plotting commands are run on processed data from these two files so the first step is to create a BlobDir dataset using the BlobToolKit `blobtools create` command. BlobToolKit v4.4.6 was used in this study and is compatible with output files from BUSCO versions 4, 5 (Manni et al. 2021) and 6 (Tegenfeldt et al. 2025) using odb10 or odb12 lineage datasets. Snail plots do not directly reveal the presence of cobionts in an assembly so this step in the workflow promotes consideration of contaminants prior to producing a snail plot and is a step that will already have been run by users running BlobToolKit as part of routine contaminant/cobiont screening (e.g. (Houliston et al. 2025 July 29)). While adding an extra step to snail plot generation, using an intermediate BlobDir has the advantage that processed data can be reused for related plots and can be filtered based on any imported variable.

Running the full BlobToolKit pipeline is computationally intensive, particularly for large assemblies, so a minimal BlobDir can be created for the purposes of generating a snail plot, if required, using a command of the form:

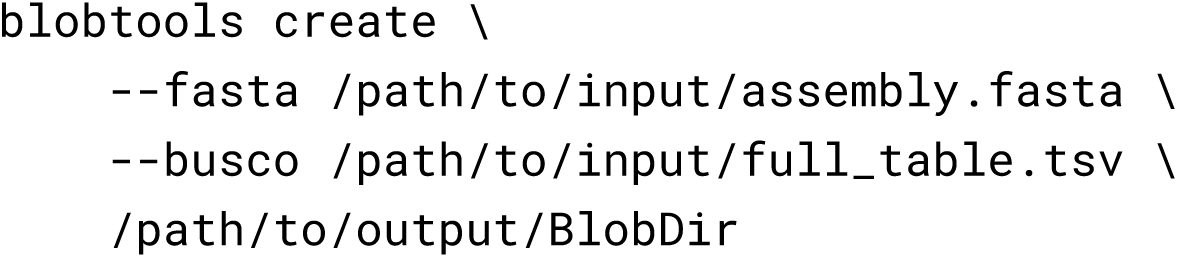

#### Plot generation

A snail plot can be generated using default settings using a command of the form:

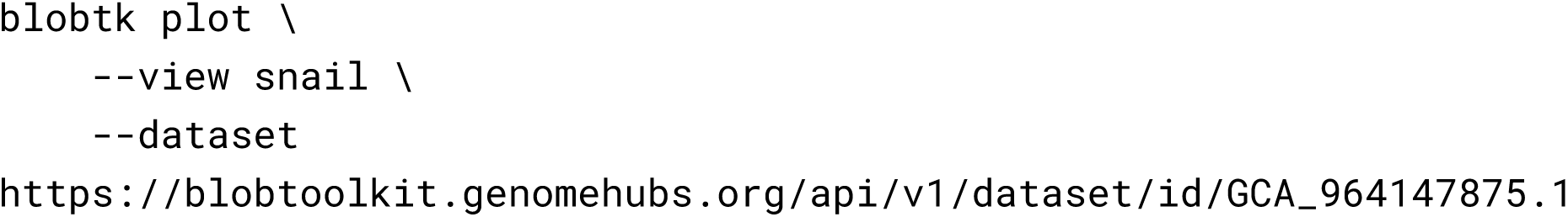

#### Plot options

Snail plots are intended to serve as assembly badges so, while there is scope for some customisation, the range of options is limited to ensure visual consistency of core plot features. Table 1 shows the available snail plot customisation options in BlobTK version 0.7.9.

**Table 1.** Snail plot customisation options in blobtk plot.

| Option | Description | Default value |
| --- | --- | --- |
| `-o, --output` | Output filename. Use .png/.svg suffix to render snail plot or .json/.yaml suffix to print all plot statistics to file | output.svg |
| `-f, --filter` | Apply a filter to exclude scaffolds based on any variable in the BlobDir dataset |  |
| `-s, --segments` | Segment count | 1000 |
| `--max-span` | Maximum value on the circumferential axis | assembly span |
| `--max-scaffold` | Maximum value on the radial axis | longest scaffold length |
| `--scale-function` | Scale function for the radial axis (linear, sqrt or log) | sqrt |
| `--significant-digits` | Significant digits to use when rounding numbers for display | 3 |
| `--decimal-precision` | Number of decimal places) to show when rounding percentages for display | 2 |
| `--rounding` | Rounding strategy (round, down or up) | round |
| `--show-numbers` | Flag to show absolute values instead of percentages | false |
| `--busco-numbers` | Flag to show absolute values instead of percentages for BUSCO counts only | false |
| `--badge` | Flag to show snail plot as an assembly badge with no legend or text | false |
| `--show-score` | Flag to show the snail score | false |

Setting the output filename sets the image format according to the filename suffix (png or svg) or allows for printing the processed plot data to a json or yaml file.

A filter option restricts the list of scaffolds included in the plot. This is intended to be used in conjunction with the interactive BlobToolKit viewer on datasets processed using the full BlobToolKit pipeline, but can also be used to see the impact of, for example, excluding contigs of less than 1kb. The syntax for filtering is consistent with that used in the BlobToolKit viewer URL and blobtools command line tool so such a filter would be specified as `--filter length--Min=1000`.

Snail plots are visually circular, but this is implemented as a polygon with a default of 1,000 sides. Within each segment, the GC and N values are summarised across all scaffolds in that bin. Reducing the number of segments to 100 increases the bin size for these summary statistics, which can reduce the noise in the GC track for portions of the assembly with large numbers of short contigs. Smaller values could be used to produce stylised plots at the expense of accuracy in representing proportional scaffold lengths..

Plots are intended to be scale-independent to allow assessment of assemblies regardless of the total assembly span or longest scaffold length. When comparing assemblies directly, the max-span and max-scaffold lengths can be set manually to produce plots with the same circumferential and radial ranges.

Snail plots were introduced before chromosome-level assemblies became commonplace so scaffold lengths are square-root scaled by default to increase visibility of trends in sequence length among the shortest scaffolds in the assembly. In assemblies with low contiguity these short scaffolds can represent a high proportion of the data.

However for chromosomal assemblies this emphasis on the shorter scaffolds reduces the resolution of differences in chromosome lengths. To address this, the scale-factor can be set to linear.

The remaining options (Table 1) all concern the display and formatting of the plot legend and axis tick labels. These allow control of the rounding behaviour and significant digits/precision separately for absolute values and percentages and displaying absolute values instead of percentages. These options don’t affect the visual consistency of the core plot features, but allow for avoiding issues with rounding to the same value when comparing assemblies with similar BUSCO completeness in lineages with large numbers of BUSCO genes. Snail plots are intended to convey more nuanced information than individual assembly metrics such as N50 can provide without reference to the underlying values. The `--badge` option fully realises the potential of snail plots to act as assembly badges by removing all legends and numerical annotation from the plot.

### Interpreting snail plots

Snail plots layer a suite of assembly metrics into a dense visual representation of assembly quality, the core plot components of which are dissected in Figure 1. Most graphical information is presented in a circular plot, where the circumferential axis is labeled with cumulative span and percentage of the total assembly (Figure 1A). Data presented along this axis are derived from size-sorted scaffolds sampled at regular intervals along the assembly span, with the default being to segment the assembly into 1,000 bins (i.e. each bin represents 0.1% of the assembly span). If a scaffold spans multiple bins, the values derived from that scaffold contribute to each bin it covers, either fully or partially. If multiple scaffolds are present in a single bin, values are summarised across all scaffolds in the bin. The outer portion of the plot contains a base composition track with dark and light blue shading representing the GC and AT proportions, respectively (Figure 1B). If scaffolds in a bin contain Ns, the proportion of Ns is represented by white shading at the edges of this track. Variation in GC content across multiple scaffolds in a bin is represented by lighter and darker shading to show the range of values about the mean. At the center of the plot, a region of light grey shading indicates the cumulative scaffold count (Figure 1C). The distance from the center of the plot for the scaffold counts is log-scaled with solid white gridlines shown at order-of-magnitude intervals from 10 up. As noted, the log scaling of this axis can be substituted with linear scaling for chromosomally-complete assemblies. The lengths of the size-sorted scaffolds are shown in dark grey shading from the circumference to the center of the plot (Figure 1D). The radial axis is scaled to the length of the longest contig in the assembly and where multiple scaffolds are present in a bin, the shading represents the length of the shortest scaffold in that bin. The longest scaffold is highlighted with red shading (Figure 1E). For assemblies with multiple scaffolds in the first bin, no red shading will be present, though the radial axis still represents the longest scaffold in the assembly. Dark and light orange shading represent the N50 and N90 scaffold lengths (Figures 1F and 1G), respectively. These shaded areas are always present, however they may not be visible for assemblies with low contiguity where the longest scaffold is orders of magnitude longer than the N50 and/or N90 value. Outside the main plot area, a smaller circular plot represents the BUSCO completeness of the assembly (Figure 1H). The shaded region represents the percentage of complete BUSCOs in the assembly, darker shading shows the percentage of duplicated BUSCOs and lighter shading shows the percentage fragmented.

**Figure 1.**
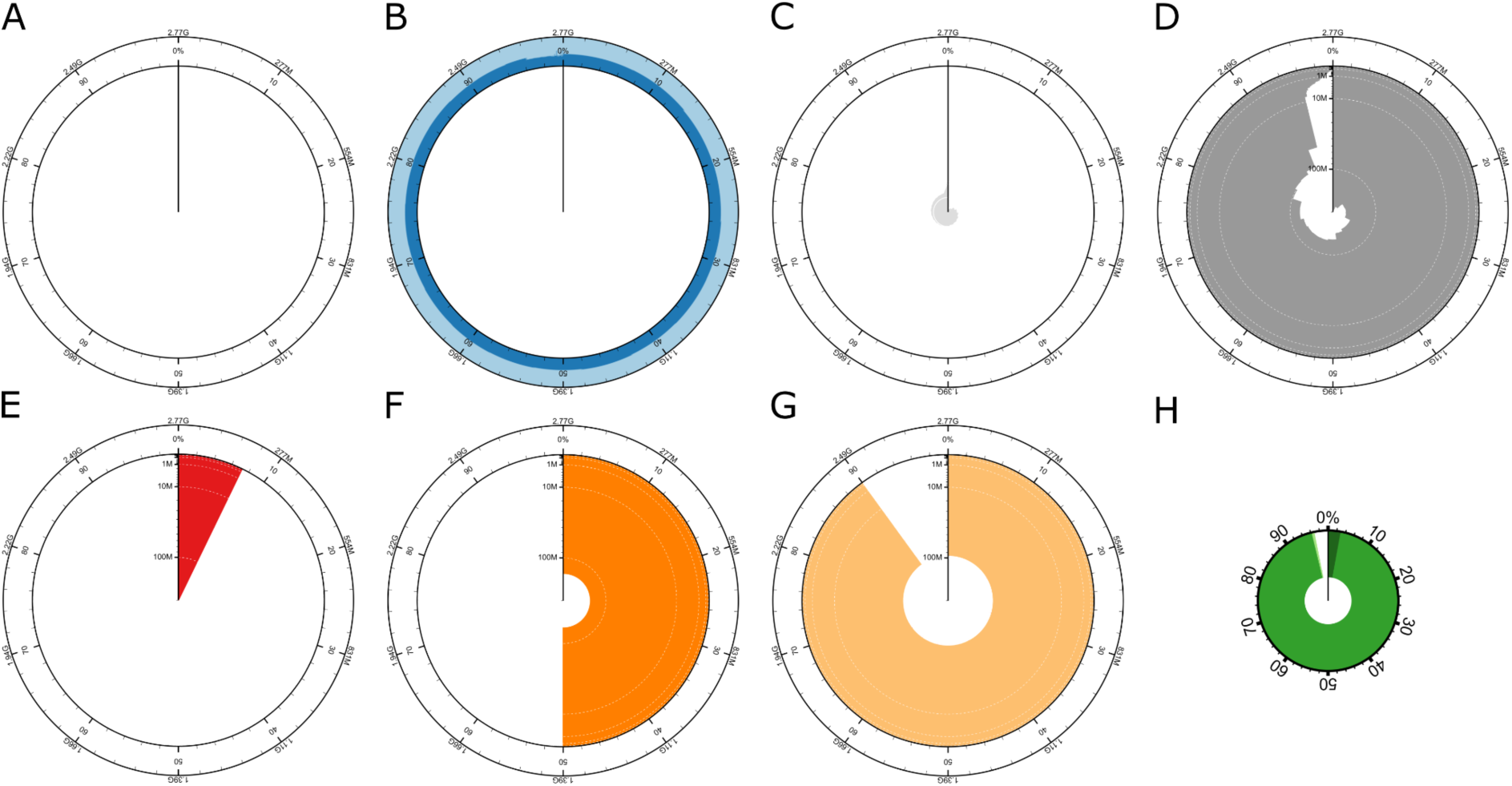
Snail plot features taken from a plot of *Mus musculus* assembly GCA_949316315.1. (A) Circumferential axis showing cumulative and percentage span, default settings divide this into 1,000 bins of 0.1% assembly span each; (B) base composition track highlighting GC and AT proportion and showing proportion of Ns if present. (C) Central light grey shading showing log-scaled cumulative count; (D) Dark grey shading showing size-sorted scaffold lengths in each bin; (E) Red shading highlights the proportional size of the longest scaffold in the assembly; (F) Dark orange shading highlights the N50 scaffold length; (G) Light orange shading highlights the N90 scaffold length; (H) an inset plot show assembly BUSCO complete, duplicated, fragmented and missing percentages. The full snail plot for this assembly is shown in Figure 2A.

### Comparing scale options

Two assemblies of the house mouse *Mus musculus*, one chromosomal (GCA_949316315.1; BlobToolKit ID: mMusMuc1_1), sequenced for the Darwin Tree of Life project (Darwin Tree of Life Project Consortium 2022) and one scaffold (GCA_000185125.1; BlobToolKit ID: AEKR01) (Gnerre et al. 2011) were chosen to highlight differences between high and low-quality assemblies and to show the impact of the scaling options on snail plot appearance. Both of these assemblies have a BlobToolKit analysis available on the public BlobToolKit server so the snail plots were generated based on the processed BlobDir data directly from the API (https://blobtoolkit.genomehubs.org/api/v1) using commands of the form:

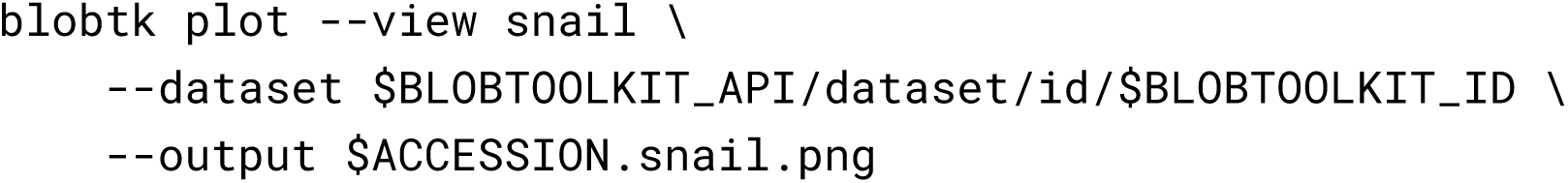

A plot was generated using the default scale function for each accession/BlobToolKit ID (Figure 2). Additional plots were generated for the low contiguity assembly GCA_000185125.1 scaled to the same total span as assembly GCA_949316315.1 using the `--max-span` option and scale to both the same total span and same maximum scaffold length by additionally specifying the `--max-scafold` option.

**Figure 2.**
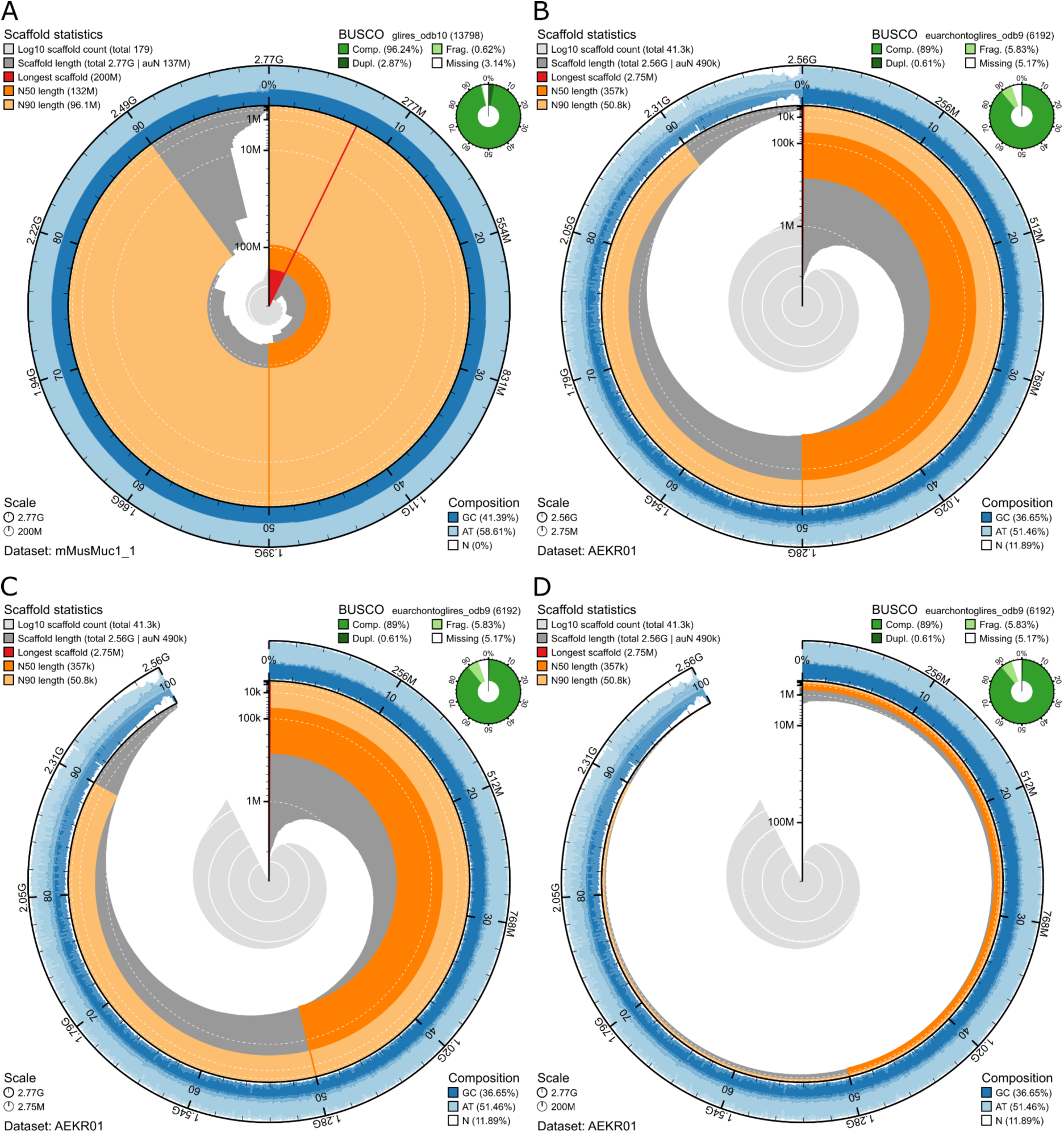
Snail plots for two assemblies of the house mouse, *Mus musculus*, highlight the visual contrast between high contiguity assembly GCA_949316315.1 and low contiguity assembly GCA_000185125.1.

Standard snail plots are shown in (A) and (B) for accessions GCA_949316315.1 and GCA_000185125.1, respectively. In (C) and (D) variants of the snail plot for assembly GCA_000185125.1 are shown scaled to the same total assembly span as GCA_949316315.1, highlighting the proportional difference in span, and in (D) the assemblies are scaled to the same longest scaffold length, emphasising the difference in N50 and N90 lengths but producing a plot with a much lower information content.

The ability of a snail plot to act as an assembly badge is dependent on consistent implementation and scaling. However, for chromosomal assemblies, coverage of the majority of the genome in a small number of scaffolds with lengths close to the longest scaffold can lead to compression of the sequence lengths on the default square-root (sqrt) scale. An option to change this to a linear scale improves the information content of the plot without hindering use as an assembly badge as the change in scale is readily apparent from the even distribution of the radial axis tick marks, labels and gridlines. Figure 3 shows the visual impact of changing the scale for the chromosomal Mus musculus assembly GCA_949316315.1. Figure 3A reproduces the snail plot from Figure 2A, but with the `--show-numbers` option to use numbers instead of percentages in the BUSCO and composition legends, This change is unlikely to be noticed at a glance, highlighting the secondary role of the numerical values in snail plot interpretation.

**Figure 3.**
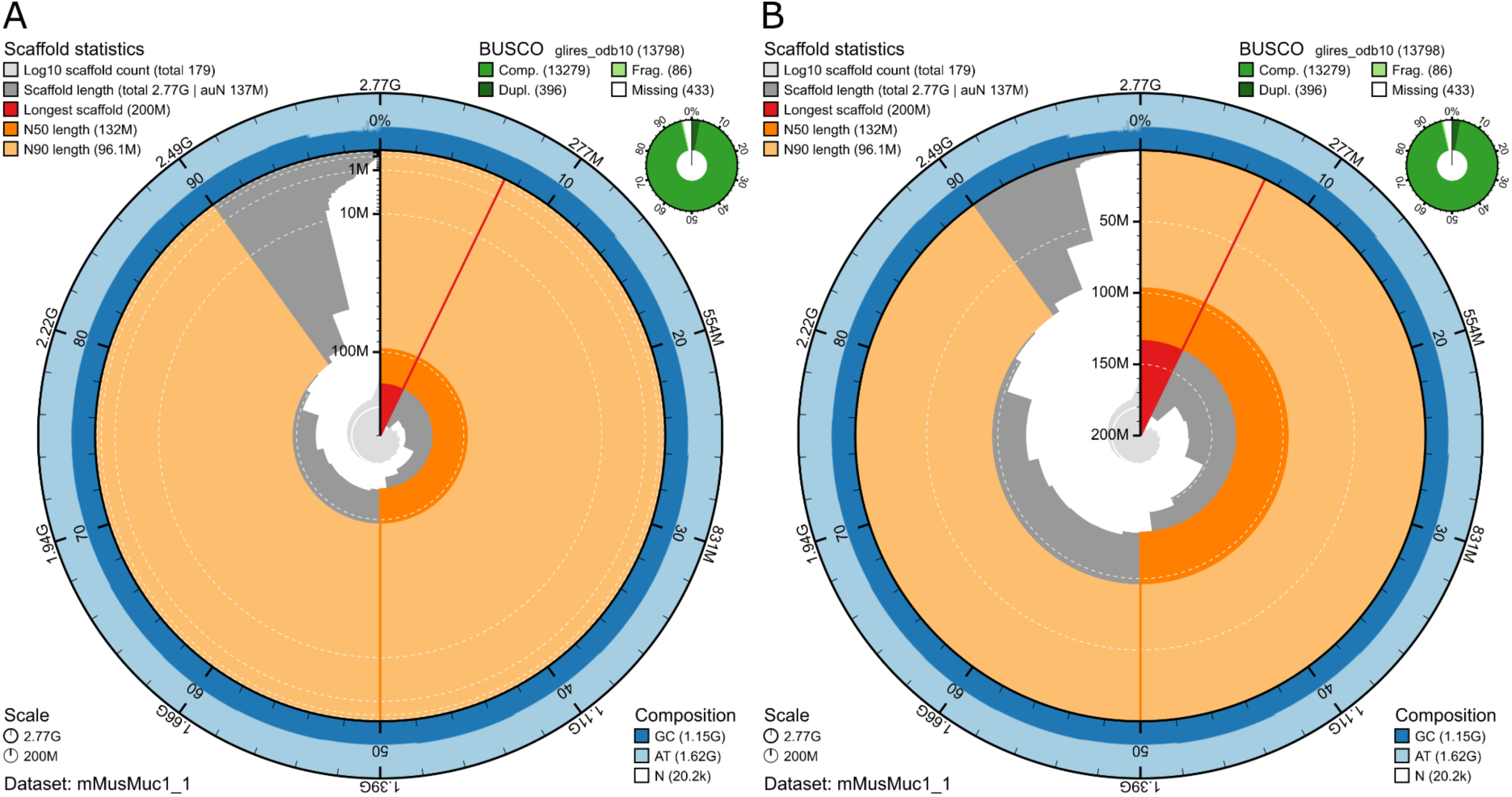
Comparison of scaffold length scale factor options on plots for the chromosomal *Mus musculus* assembly GCA_949316315.1: (A) using the default sqrt scale and (B) using the optional linear scale.

The distribution of scaffold lengths in Figure 3A demonstrates the problem of compression of information content on the sqrt scale, compared with Figure 3B which uses a linear scale. The prominent label at 200 Mb and gridlines at 150 Mb and 50 Mb help with interpretation of the physical sizes of scaffolds up to and beyond the N90 length, while the linear scale also allows direct assessment of the relative lengths of scaffolds throughout the distribution.

### Correlation with standard metrics

A set of assemblies at scaffold, chromosome or complete genome level with scaffold N50 > 10 kb, contig N50 > 1 kb, assembly span > 10 Mb and BUSCO completeness > 80% was obtained from the 2025.04.21 archive release of Genomes on a Tree (Challis et al. 2023). Assemblies were grouped into bins by order of magnitude of scaffold and contig N50 values and a single representative assembly was chosen pseudo-randomly from each bin using the python `random` module with a seed of 1031. The corresponding BlobToolKit ID was identified for each assembly and used to generate a snail plot using a `blobtk plot` command as above, with output files saved in SVG and JSON formats. SVG files for each plot were combined into a single file to generate a grid of snail plots on axes of increasing contig and scaffold N50, with one plot per occupied bin. This process was automated using the docs/snail-plots/figure4.py script in the BlobTK repository.

### Taxonomic distribution of example assemblies

A taxonomic tree view of the included assemblies was generated using the 2025.14.21 acrhival release of Genomes on a Tree. The tree was coloured by taxonomic kingdom with bars to indicate assembly span using the URL https://goat.genomehubs.org/search?query=assembly_id%3DGCA_900322205.1%2CGCA_003016195.1%2CGCA_001632505.1%2CGCA_000261425.2%2CGCA_002222395.1%2CGCA_020883555.1%2CGCA_001661245.1%2CGCA_000204055.1%2CGCA_964340765.1%2CGCA_013467465.1%2CGCA_014337955.1%2CGCA_018257905.1%2CGCA_964340405.1%2CGCA_000001215.4%2CGCA_003033685.1%2CGCA_013339765.2%2CGCA_963691655.1%2CGCA_949316315.1%2CGCA_019009955.1%2CGCA_964205295.1%2CGCA_963693085.1&result=assembly&taxonomy=ncbi&report=tree&collapseMonotypic=true&treeStyle=ring&y=assembly_span&cat=kingdom&hideSourceColors=true. The tree was manually edited to ensure all GCA accession labels were printed in full.

### Summary statistic

The area under the N*x* curve, auN (Li 2020) or E-size (Salzberg et al. 2012), was calculated by dividing the sum of squared scaffold lengths by the sum of scaffold lengths (or assembly span). By this formula, the value of auN will always be less than or equal to the length of the longest scaffold, and in practice can only be equal to the longest scaffold length if the assembly has a single scaffold or if all scaffolds are of equal length. As such, a relative measure of area under N can be derived by dividing auN by the longest scaffold length to give a proportional value <= 1 of auN relative to the longest scaffold length.

Joining relatively short contigs into scaffolds with runs of N is common practice when scaffolding assemblies. For assemblies with a high proportion of Ns, this relative auN value could be inflated by joining short contigs with long runs of N, resulting in an assembly with higher contiguity but relatively low information content compared with a similar assembly with a low proportion of Ns. To account for this aspect of quality, we also calculate a corrected auN value by substituting the square of the number of ACGT bases per scaffold into the standard auN calculation in place of scaffold length and derive a corrected relative auN value by dividing the corrected auN by the longest scaffold length, including Ns. This corrected value will be equal to relative auN for assemblies with no Ns and below relative auN for assemblies with a high proportion of Ns. Since this statistic has been developed in response to the visual characteristics of snail plots, we term the corrected relative auN value for an assembly the assembly “snail score”.

## Results & Discussion

Snail plots are intended to act as badges of assembly quality. The focus is on rapid interpretation of quality at a glance, but the full plots also include legends with values for a range of core statistics, including total assembly span, scaffold N50 and BUSCO completeness. This allows both visual and numerical inspection of the differences between two assemblies for the house mouse, *Mus musculus* (Figure 2). Consideration of the main plot features suggests that chromosomal assembly GCA_949316315.1 (Figure 2A) is of higher quality overall than scaffold assembly GCA_000185125.1 (Figure 2B).

Overall, the scaffold length curve occupies a much larger proportion of the plot area for assembly GCA_949316315.1 and has a stepped profile compared with the smooth curve for assembly GCA_000185125.1. The longest scaffold and N90 overlays are both more prominent for assembly GCA_949316315.1 and the N50 overlay extends further towards the center of the plot and its prominence is only reduced by the extent of the N90 overlay. Around the outside of the plot, assembly GCA_949316315.1 has a more consistent GC proportion compared to assembly GCA_000185125.1, which also has a higher proportion of whitespace on the composition track, indicating a higher proportion of N bases in the assembly. Assembly GCA_949316315.1 also has higher BUSCO completeness, although noting that these assemblies were processed for BlobToolKit at different times and used different BUSCO versions and lineage datasets. Table 2 presents a detailed side-by-side comparison of individual plot features for the two assemblies.

**Table 2.**
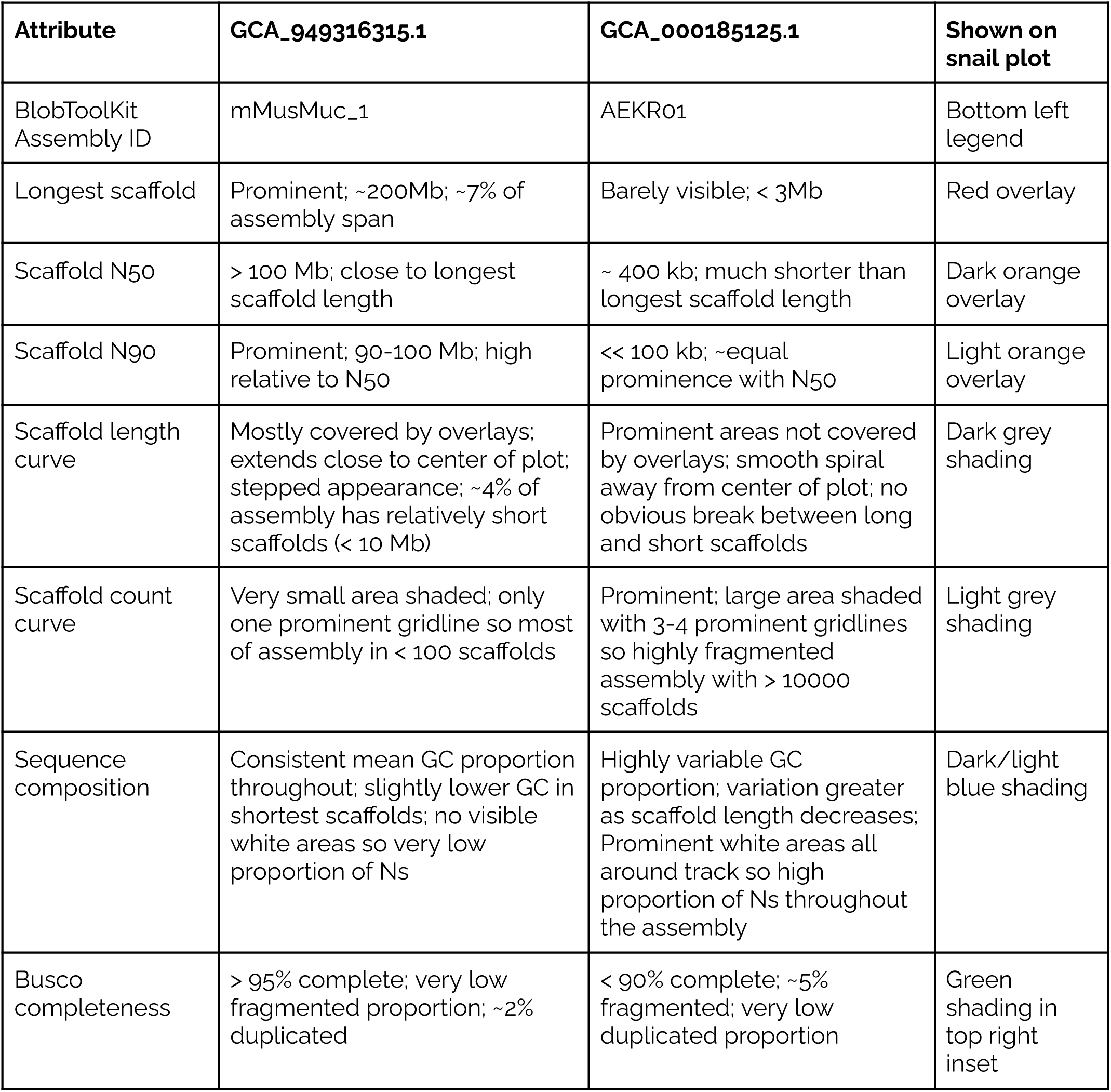
Pairwise comparison of snail plot features for *Mus musculus* assemblies GCA_949316315.1 and GCA_000185125.1 as shown in Figures 2A and 2B, respectively.

| Attribute | GCA_949316315.1 | GCA_000185125.1 | Shown on snail plot |
| --- | --- | --- | --- |
| BlobToolKit Assembly ID | mMusMuc_1 | AEKR01 | Bottom left legend |
| Longest scaffold | Prominent; ~200Mb; ~7% of assembly span | Barely visible; < 3Mb | Red overlay |
| Scaffold N50 | > 100 Mb; close to longest scaffold length | ~ 400 kb; much shorter than longest scaffold length | Dark orange overlay |
| Scaffold N90 | Prominent; 90-100 Mb; high relative to N50 | << 100 kb; ~equal prominence with N50 | Light orange overlay |
| Scaffold length curve | Mostly covered by overlays; extends close to center of plot; stepped appearance; ~4% of assembly has relatively short scaffolds (< 10 Mb) | Prominent areas not covered by overlays; smooth spiral away from center of plot; no obvious break between long and short scaffolds | Dark grey shading |
| Scaffold count curve | Very small area shaded; only one prominent gridline so most of assembly in < 100 scaffolds | Prominent; large area shaded with 3-4 prominent gridlines so highly fragmented assembly with > 10000 scaffolds | Light grey shading |
| Sequence composition | Consistent mean GC proportion throughout; slightly lower GC in shortest scaffolds; no visible white areas so very low proportion of Ns | Highly variable GC proportion; variation greater as scaffold length decreases; Prominent white areas all around track so high proportion of Ns throughout the assembly | Dark/light blue shading |
| Busco completeness | > 95% complete; very low fragmented proportion; ~2% duplicated | < 90% complete; ~5% fragmented; very low duplicated proportion | Green shading in top right inset |

Many of these key differences are also apparent from inspection of the set of numerical values in the plot legends that would commonly be presented in a table. The chromosomal assembly GCA_949316315.1 has longest scaffold, N50 and N90 lengths all orders of magnitude greater than the equivalent values for assembly GCA_000185125.1. There are also large differences in the number of scaffolds and percentage of missing data with the ∼41.3k scaffolds of assembly scaffolds of assembly GCA_000185125.1 having an average 11.89% N bases compared to 0% N bases for the 179 scaffolds of assembly GCA_949316315.1. One key difference that is apparent from the values but not the standard snail plot presentation is that assembly GCA_949316315.1 is 210 Mb longer. Given that these are assemblies of the same species, it would be valid to rescale the plot for assembly GCA_000185125.1 to match the greater assembly span of the chromosomal assembly (Figure 2C), which highlights the proportional difference between the total span of the two assemblies. The radial axis may also be rescaled to match (FIgure 2D), however with such a large difference in longest scaffold length, this generates a plot with mostly white space. While this may serve to highlight the difference in assembly quality, it does so at the expense of providing insight into the distribution of scaffold lengths in the less contiguous assembly such that a rescaled plot no longer serves as an assembly badge.

Arranging a set of snail plot assembly badges for assemblies of varying quality along axes of contig and scaffold N50 (Figure 4) reveals some correlation with indicators of assembly quality but highlights further assembly features that are not directly related to these primary axes. Details of the assemblies included in this plot are shown in Table 3. Individual assemblies in this figure are referred to using the Vertebrate Genome Project convention of referring to an assembly by the order-of-magnitude of the Contig and scaffold N50 lengths, respectively (Rhie et al. 2021), e.g. the last assembly on the bottom row of the plot with a 10^3^ contig N50 and and a 10^7^ scaffold N50 would be referred to as the assembly at 3.7.

**Figure 4.**
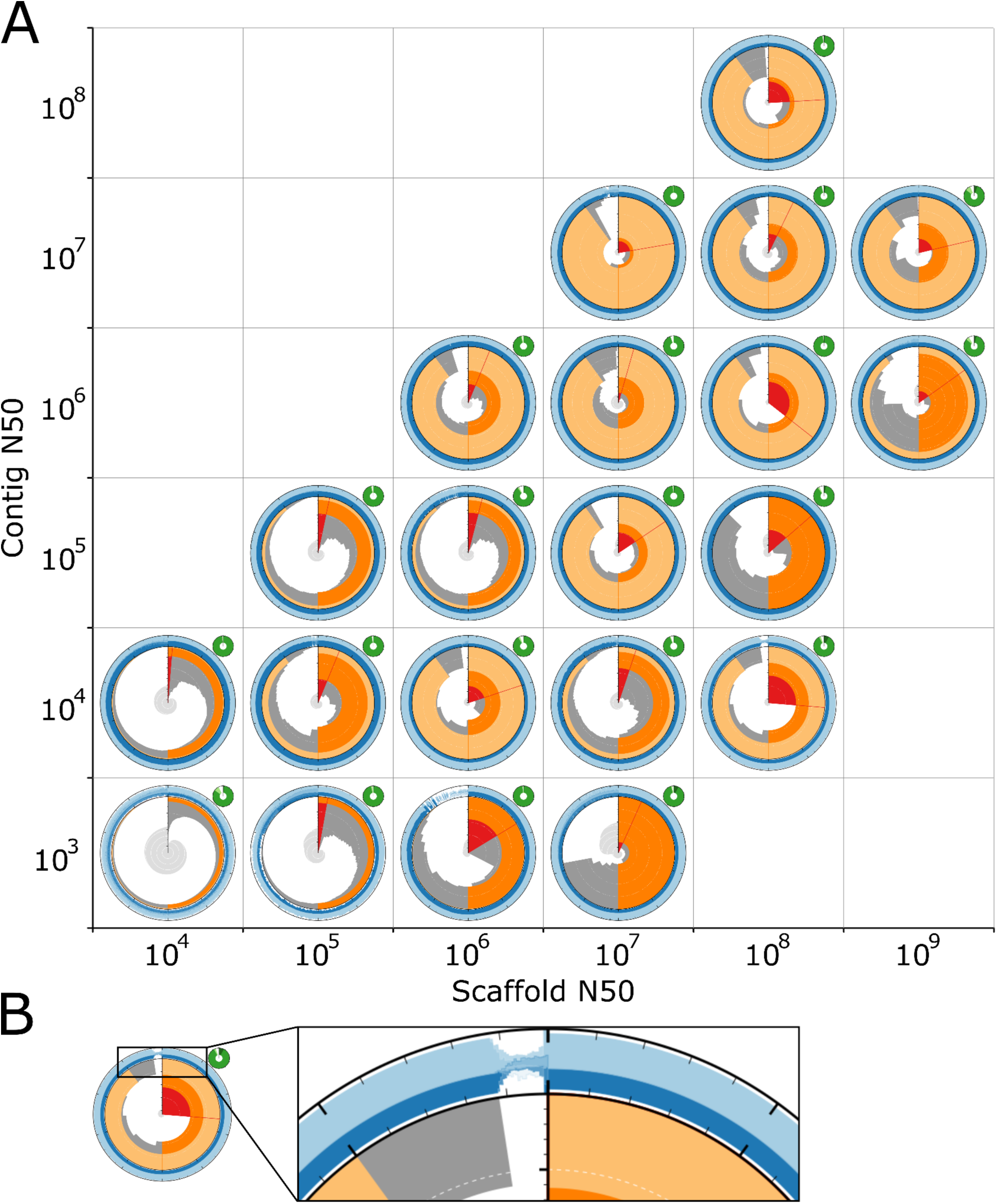
(A) A set of linear-scaled snail plots for assemblies of varying quality arranged by order of magnitude of assembly contig and scaffold N50 values. Plots are presented as assembly badges to highlight the suitability of the plots for qualitative assessment of assemblies. While features such as the amount of data ink (proportional to auN statistics) are broadly correlated with scaffold and contig N50, there are a number of differences that are less closely correlated with variation along these axes. (B) enlarged section of *Nelumbo nucifera* assembly GCA_003033685.1 (contig N50 = 10^4^, scaffold N50 = 10^8^) to show the presence of whitespace, indicating Ns in the longest and shortest scaffolds.

**Table 3.**
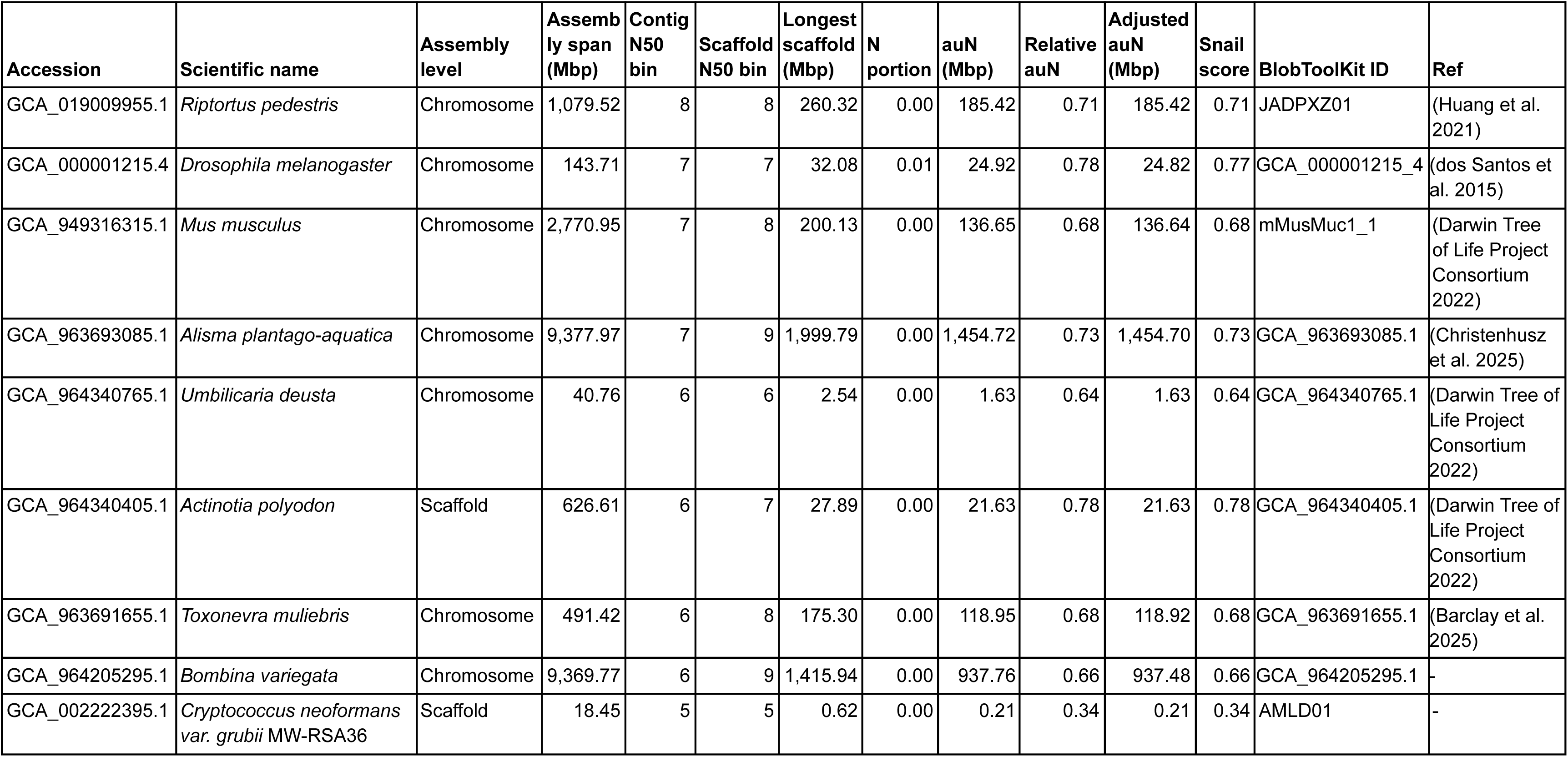
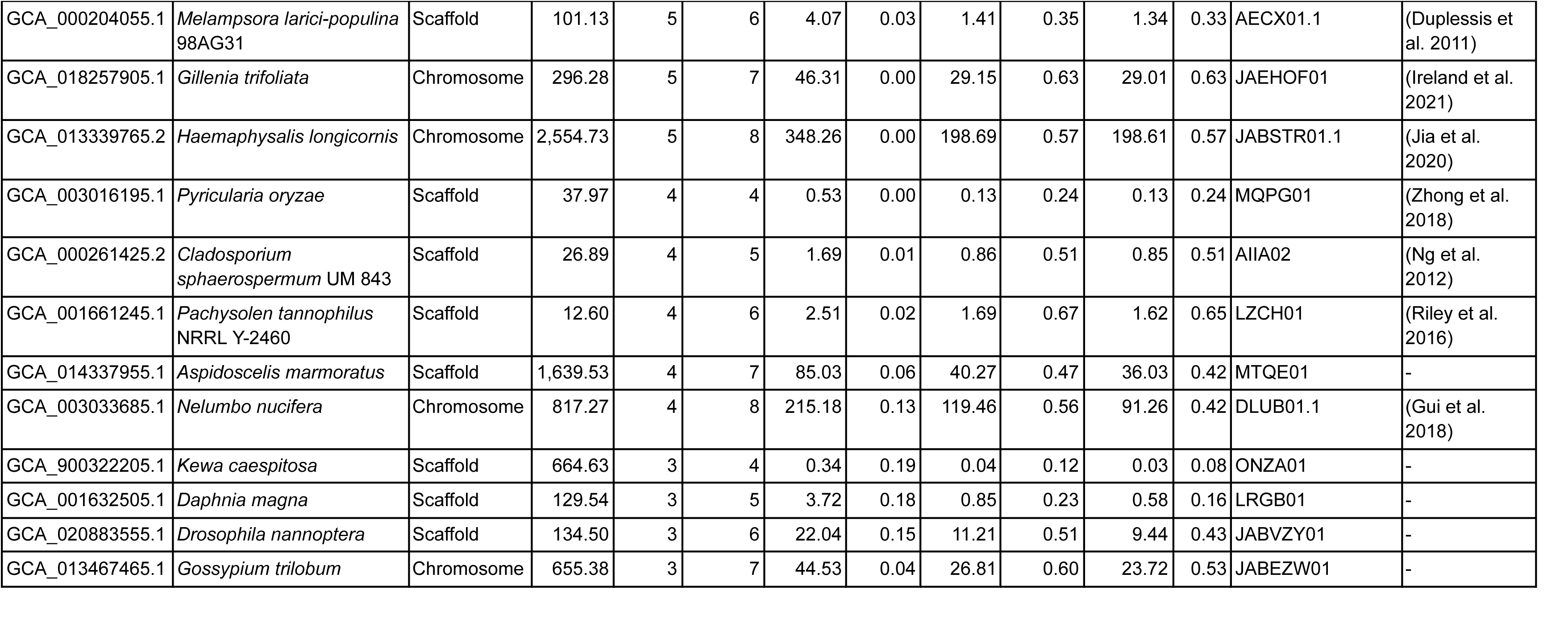
Selected assembly statistics for assemblies included in the assessment of correlation between N50 values and snail plot assembly badge features.

Considering the key features highlighted above, assemblies towards the top right of the plot appear to have higher quality than those at the bottom left, sharing prominent N50 and N90 overlays, a stepped scaffold length curve that fills much of the plot area and a small scaffold count curve. By contrast assemblies towards the bottom left have prominent scaffold count curves, smooth scaffold length curves and less prominent N50 and in particular N90 overlays. Among assemblies with relatively high N90, there is a distinction between those for which scaffold lengths in each bin are within an order of magnitude of the longest scaffold length and those with a large number of relatively short scaffolds showing as a gap in the circularised scaffold length distribution. For *Drosophila melanogaster* assembly GCA_000001215.4 at 7.7, this is accompanied by an increase in sequence composition variability and/or proportion of Ns.

Some assemblies with low contig N50 (in the 10^3^ and 10^4^ bp bins) have scaffold N50 several orders of magnitude greater than contig N50. These typically show signatures of low quality assemblies as discussed above, but the assemblies at 4.6 and 4.8 (*Pachysolen tannophilus* NRRL Y-2460 assembly GCA_001661245.1 and *Nelumbo nucifera* assembly GCA_003033685.1, respectively) have several signatures of high quality assemblies. A key difference between these assemblies is that assembly GCA_001661245.1 at 4.6 has an almost completely shaded sequence composition track, reflecting an overall proportion of Ns < 2%, which assembly GCA_003033685.1 at 4.8 has whitespace at the edges all around this track, reflecting > 13% Ns across the whole assembly (see Figure 4B). Assembly GCA_001661245.1 could therefore be judged to be of higher overall quality.

Placing the assemblies selected for the comparison above on a taxonomy tree using GoaT shows that the assemblies span the tree of life and have a range of assembly spans from 12.6 Mb to 9.4 Gb (Figure 5). This demonstrates the effectiveness of snail plots for providing insight into markers of assembly quality across three kingdoms in the Eukaryota and across almost 4 orders of magnitude, emphasising the scale-independence of the plots.

**Figure 5.**
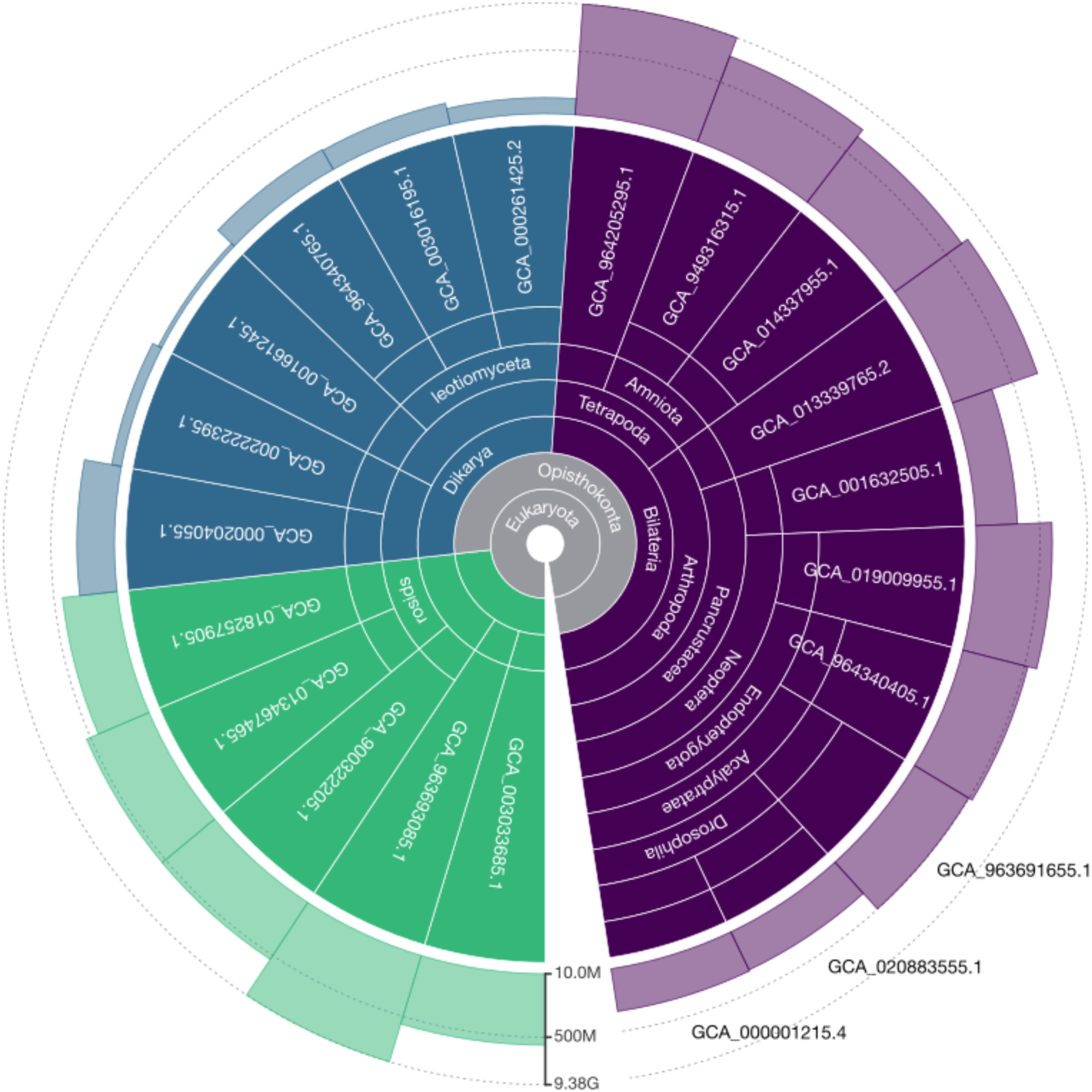
Taxonomic distribution and sizes of assemblies included in the assessment of correlation between N50 values and snail plot assembly badge features. Taxa are coloured by kingdom to highlight plant (green), fungus (blue) and animal (purple) genomes. Bars show the span of each assembly ranging from 12.6Mb to 9.38Gb. Report generated using <u>Genomes on a Tree</u>.

The interpretation of quality above emphasises the importance of the area covered by the scaffold length curve, including the associated overlays. This is closely related to the area under N (auN), however, auN lacks the scale-independence of snail plots since the maximum possible value of auN increases with assembly span. To correct for this, we propose using a relative auN score by dividing auN by the longest scaffold length to obtain a score between 0 and 1. The relative auN score reflects the scaling used when drawing snail plots and scores for the assemblies in FIgure 4 are shown in Table 3.

To account for the proportion of ambiguous bases (Ns), another key indicator of quality, an adjusted variant of the relative auN score can be used. The adjustment is based on squaring the count of ACGT bases in each scaffold when calculating auN instead of simply squaring the scaffold lengths. There is a good correlation between snail plot features associated with higher assembly quality and this corrected value, which we term the assembly “snail score”. While this is necessarily a somewhat subjective correlation, as shown in Figure 6, snail scores broadly increase with contig and scaffold N50. The highest snail scores of 0.77 and 0.78 are obtained by assemblies at 7.7 and 6.7, *Drosophila melanogaster* assembly GCA_000001215.4 and *Actinotia polyodon* assembly GCA_964340405.1, respectively. Among the assemblies with contig N50 values below 10^6^, the two assemblies with the most visual indicators of high assembly quality are the only assemblies with snail scores above 0.6, these are *Gillenia trifoliata* assembly GCA_018257905.1 at 5.7 with a score of 0.63 and *Pachysolen tannophilus* NRRL Y-2460 assembly GCA_001661245.1 at 4.6 with a score of 0.65. *Nelumbo nucifera* assembly GCA_003033685.1 at 4.8, which was contrasted with GCA_001661245.1 above as having visual markers of high assembly quality but a high proportion of Ns achieves an unadjusted (relative auN) score of 0.56, but an adjusted, snail score of 0.42.

**Figure 6.**
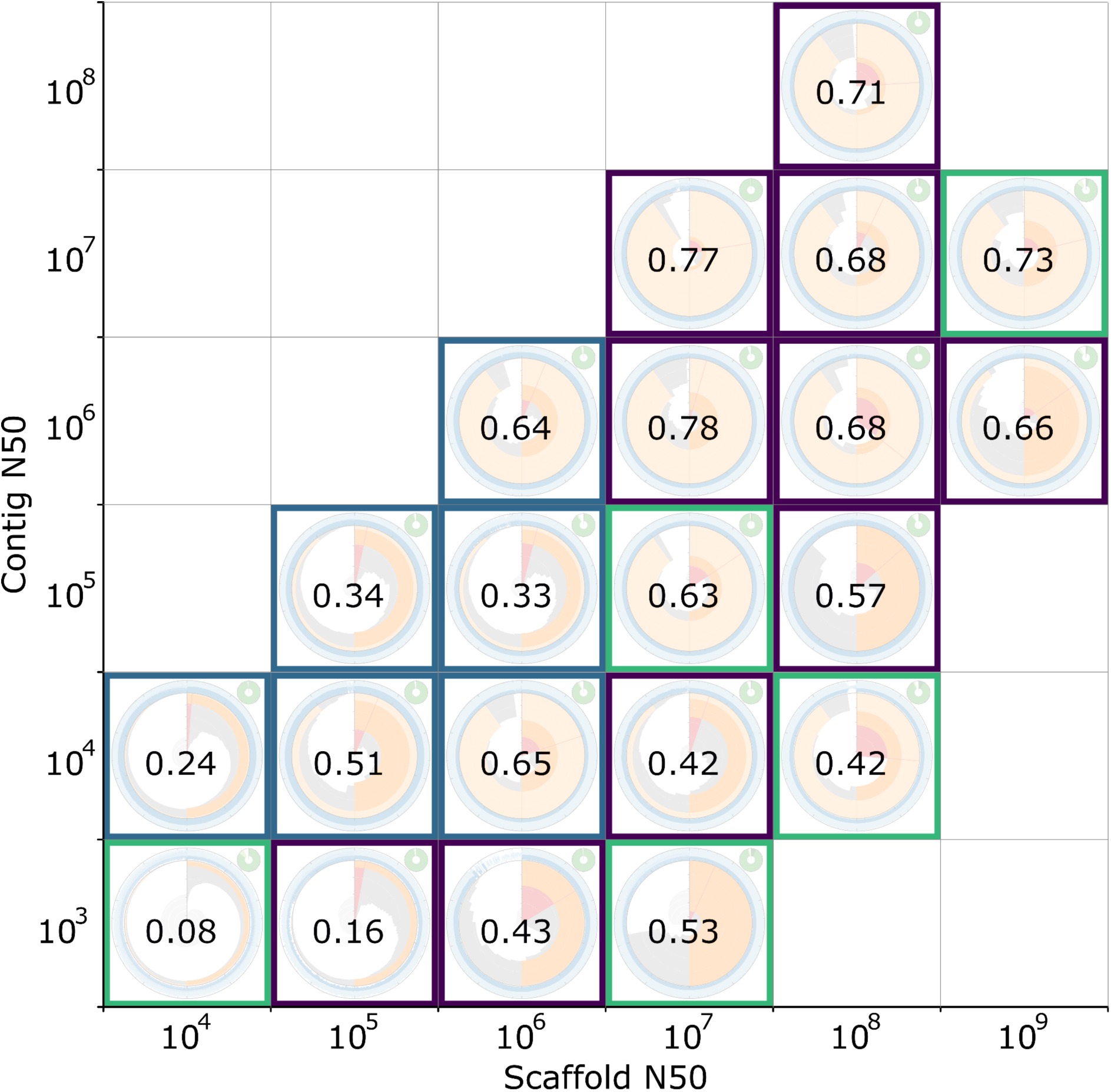
Snail scores (relative measure of auN adjusted for ambiguous bases) for the assemblies in Figure 3. Coloured borders indicate the taxonomic kingdom, plant (green), fungus (blue) and animal (purple), for each assembly.

Based on this sample of assemblies, it is apparent that snail scores correlate with visual markers of assembly quality and, while acknowledging that any cutoff is necessarily arbitrary, we suggest that a score above ∼0.6 can be taken as a broad indication of assembly quality. Assemblies within the sample set that achieved a score above this cutoff were drawn from three kingdoms of Eukaryota and had a range of assembly spans from 12.6 Mb to 9.4 Gb. Thus, snail scores provide a single value indicator of assembly quality, with higher values associated with more markers of assembly quality, allowing them to be used for assessments of relative assembly quality across a wide range of taxonomic and assembly size diversity. It should be noted, however, that the maximum possible snail score for a taxon is limited by the relative physical sizes of its chromosomes. Taxa with a single large chromosome and a number of much smaller chromosomes will have relatively low snail scores, even for a full telomere-to-telomere assembly.

## Conclusions

Snail plots have become a widely used representation of core genome assembly metrics that facilitate rapid identification of characteristic markers of both high and low assembly quality. These markers of quality are broadly correlated with assembly N50 values, and perhaps most closely with metrics that consider the full distribution of N*x* values. Unlike this family of metrics, snail plots present this information in a scale-independent way, similar to plots of the full N*x* curve. Based on this observation, we have suggested modifying the area under N (auN) calculation to account for ambiguous bases and scale to a proportion of the maximum auN score possible given the assembly span and longest scaffold length. The resulting snail score correlates closely with a visual assessment of assembly quality markers in a set of snail plots. While snail scores could fulfill the requirement for a single tabulatable value to indicate assembly quality across a range of genome sizes, snail plots are still a valuable tool for a more holistic view of assembly quality.

## Data availability

The `blobtk plot` command is available under an MIT license from the BlobTK GitHub repository at https://github.com/genomehubs/blobtk. This repository also contains scripts and commands used to generate figures presented in this study under the docs/snail-plots directory. Version 0.7.9 used in this study is archived at https://zenodo.org/records/17661559. All data analysed are publicly available via https://goat.genomehubs.org/2025.04.21/ and from https://blobtoolkit.genomehubs.org.

## Acknowledgments

Snail plots have been improved by community engagement with the current and previous implementations. We are particularly grateful to users who submitted bug reports and feature requests to our GitHub repositories and we thank Amy and Ayla Challis for comments on drafts of this manuscript. Portions of the snail plot code in BlobTK were developed with GitHub Copilot AI autocompletion enabled.

## Study funding

This research was funded in whole, or in part, by the Wellcome Trust 220540/Z/20/A. For the purpose of Open Access, the author has applied a CC BY public copyright licence to any Author Accepted Manuscript version arising from this submission.

